# The Neural Response at the Fundamental Frequency of Speech is Modulated by Word-level Acoustic and Linguistic Information

**DOI:** 10.1101/2022.04.08.487621

**Authors:** Mikolaj Kegler, Hugo Weissbart, Tobias Reichenbach

## Abstract

Spoken language comprehension requires rapid and continuous integration of information, from lower-level acoustic to higher-level linguistic features. Much of this processing occurs in the cerebral cortex. Its neural activity exhibits, for instance, correlates of predictive processing, emerging at delays of a few hundred milliseconds. However, the auditory pathways are also characterized by extensive feedback loops from higher-level cortical areas to lower-level ones as well as to subcortical structures. Early neural activity can therefore be influenced by higher-level cognitive processes, but it remains unclear whether such feedback contributes to linguistic processing. Here, we investigated early speech-evoked neural activity that emerges at the fundamental frequency. We analyzed EEG recordings obtained when subjects listened to a story read by a single speaker. We identified a response tracking the speaker’s fundamental frequency that occurred at a delay of 11 ms, while another response elicited by the high-frequency modulation of the envelope of higher harmonics exhibited a larger magnitude and longer latency of about 18 ms. Subsequently, we determined the magnitude of these early neural responses for each individual word in the story. We then quantified the context-independent frequency of each word and used a language model to compute context-dependent word surprisal and precision. The word surprisal represented how predictable a word is, given the previous context, and the word precision reflected the confidence about predicting the next word from the past context. We found that the word-level neural responses at the fundamental frequency were predominantly influenced by the acoustic features: the average fundamental frequency and its variability. Amongst the linguistic features, only context-independent word frequency showed a weak but significant modulation of the neural response to the high-frequency envelope modulation. Our results show that the early neural response at the fundamental frequency is already influenced by acoustic as well as linguistic information, suggesting top-down modulation of this neural response.

## 1 INTRODUCTION

Spoken language consists of both lower-level acoustic as well as higher-level linguistic information that need to be rapidly and continuously processed in the brain (Giraud and Poeppel (2012); Meyer (2018); Brodbeck and Simon (2020)). The lower level acoustic processing is thereby typically attributed to the primary auditory cortex, and the processing of higher-level information to the secondary auditory cortex as well as other cortical areas such as the prefrontal cortex (Hickok and Poeppel (2007); Peelle et al. (2010); Golumbic et al. (2012)).

Linguistic processing encompasses both context-independent and context-dependent aspects. An important context-independent aspect is word frequency, that is, the frequency of a word in a large text corpus (Baayen (2001)). This information has been found to be reflected in neural activity from the cerebral cortex (Brennan et al. (2012, 2016); Brodbeck et al. (2018)). Context-dependent processing is another important linguistic aspect of speech encoding, especially in noisy auditory scenes. Behavioural studies have, for instance, shown that sentences with missing parts or added noisy intrusions can still be understood by the participants (Miller and Isard (1963); Warren (1970); Rubin (1976); Dilley and Pitt (2010); Clarke et al. (2014)).

The word expectancy resulting from context is reflected in cortical responses. Indeed, words elicit a cortical negativity at a latency of about 400 ms, the N400 response, and the N400 is modulated by word expectancy (Kutas and Hillyard (1984)). Word prediction and violations of such predictions are reflected in further aspects of cortical activity such as the beta- and gamma-band power, as has been found in studies using single sentences (Friederici et al. (1993); Friederici (2002); Bastiaansen and Hagoort (2006); Kutas and Federmeier (2011); Kielar et al. (2014)). Moreover, we and others recently showed that cortical activity recorded from electroencephalography (EEG) acquired when subjects listened to stories consisting of many sentences exhibited correlates of word surprisal, that is, of the violation of word predictions, as well as of the precision at which predictions were made (Donhauser and Baillet (2020); Weissbart et al. (2020); Gillis et al. (2021)). The word-level surprisal is thereby defined as the conditional probability of the current word, given previous words. The word-level precision is the inverse entropy of the word, given the past context. Cortical activity during natural story comprehension has also been found to reflect the semantic dissimilarity between consecutive words (Broderick et al. (2018, 2019)).

Although the auditory system is often viewed as a feed-forward network of different neural processing stages, there exist corticofugal feedback connections from the cortex to the midbrain as well as to different parts of the auditory brainstem (Huffman and Henson Jr (1990); Winer (2005)). A particular early neural response to speech that can potentially be under such top-down control is the neural tracking of the fundamental frequency. Voiced parts of speech are characterized by a fundamental frequency, typically between 100 Hz and 300 Hz, as well as many higher harmonics. The elicited neural activity as recorded by EEG exhibits a response primarily at the fundamental frequency, as well as, to a lesser extent, at the higher harmonics (Skoe and Kraus (2010); Chandrasekaran and Kraus (2010)). The response has a short latency of around 10 ms and originates mainly in the auditory brainstem and in the midbrain, although cortical contributions have been discovered recently as well (Chandrasekaran and Kraus (2010); Bidelman (2018); Coffey et al. (2016, 2017, 2019)).

The early neural response at the fundamental frequency of speech can reflect different aspects of speech processing. It can, in particular, be shaped by language experience as well as by musical training (Wong et al. (2007); Krishnan et al. (2010); Bidelman et al. (2011); Kraus et al. (2017)). In addition, we recently showed that this response is modulated by selective attention to one of two competing speakers (Forte et al. (2017); Etard et al. (2019)). Moreover, a strong bidirectional functional connectivity between cortical and subcortical areas through corticofugal pathways was found in a speech-in-noise perception task (Price and Bidelman (2021)).

The frequency-following response (FFR) to the frequency of a pure tone can occur in a similar frequency range as the neural response at the fundamental frequency of speech, and presumably reflects related processing. This FFR may be under top-down control as well. An oddball paradigm in which many repeated tones are presented together with occasional deviant tones showed that the FFR is larger for expected than for unexpected ones, although a later study could not replicate the effect (Slabu et al. (2012); Font-Alaminos et al. (2021)). Invasive recordings in animals likewise showed correlates of prediction errors at different subcortical as well as cortical stages (Parras et al. (2017)).

Whether the early neural response at the fundamental frequency of speech is modulated by linguistic processing has not yet been investigated. A main difficulty is thereby the complexity of natural speech that complicates both the measurement of the neural response and the assessment of its modulation through linguistic information. However, recent studies have developed the methodology to measure the neural response at the fundamental frequency of speech even for continuous, non-repetitive speech stimuli. We recently proposed an approach in which we extracted a fundamental waveform from voiced speech, that is, a waveform that, at each time instance, oscillated at the time-varying fundamental frequency of speech (Forte et al. (2017)). We then related this waveform to EEG that was recorded simultaneously through linear regression with regularization (Etard et al. (2019)). Kulasingham et al. (2020) recently used a similar method to show that high-frequency modulation of the envelope is also tracked by the early neural response at the fundamental frequency.

Here, we employed the recently developed methodology to measure neural responses at the fundamental frequency of individual words that occur in continuous natural speech. We also quantified key word features, including both acoustic and linguistic ones. We then investigated whether the early response to speech was shaped by these word-level features.

## 2 MATERIALS AND METHODS

### 2.1 Dataset

We analyzed EEG responses to continuous speech that were collected for an earlier study on cortical correlates of word prediction (Weissbart et al. (2020)). The recording of this dataset is described in detail below.

### 2.2 Participants

13 young and healthy native English speakers (25 ± 3 years, 6 females) were recruited for the experiment. They were all right-handed and had no history of hearing or neurological impairment. All volunteers provided written informed consent. The experimental protocol was approved by the Imperial College Research Ethics Committee.

### 2.3 Experimental setup

The experiment consisted of a single session of EEG recording. During the experiment, the participants listened to continuous narratives in the form of audiobooks that were openly available at ‘*librivox.org*’. In particular, we used three short stories: *‘Gilray’s flower pot’, ‘My brother Henry*’ by J.M. Barrie and *‘An undergraduate’s aunt*’ by F. Anstey Patten (1910)^1^. Both audiobooks were read by a male speaker, Gilles G. Le Blanc. The total length of the audio material was 40 min. The stories were presented in 15 parts, each approximately 2.6 min long (2.6 min ± 0.43 min). The acoustic signals were presented to the participants through Etymotic ER-3C insert earphones (Etymotic, USA) at 70 dB SPL. The audiobooks’ transcriptions used for computing word-level features were obtained from the project Gutenberg^2^.

After each part, the participants answered multiple-choice comprehension questions presented on a monitor. Each participant was asked 30 questions throughout the experiment. The questions were designed to keep the volunteers engaged and to assess whether they paid attention to the stories. The participants answered the questions with an average accuracy of 96%, showing that they paid attention to the audio material and understood it.

### 2.4 EEG acquisition

The brain activity of the participants was measured using a 64-channel EEG system (active electrodes, actiCAP, and EEG amplifier actiCHamp, BrainProducts, Germany). The left ear lobe served as a reference. The impedance of all EEG electrodes was kept below 10 kΩ. The audio material presented to the participants was simultaneously recorded through an acoustic adapter (Acoustical Stimulator Adapter and StimTrak, BrainProducts, Germany) and used for aligning the EEG recordings to the audio signals. The EEG and the audio data were both recorded at a sampling rate of 1 kHz.

### 2.5 Auditory stimulus representations

For modelling the early neural response at the fundamental frequency of speech from the high-density EEG, we followed the methodology developed in Etard et al. (2019). In particular, we used the fundamental waveform as well as the high-frequency envelope modulation extracted from the speech signals as the audio stimulus features (Fig. 1).

**Figure 1.**
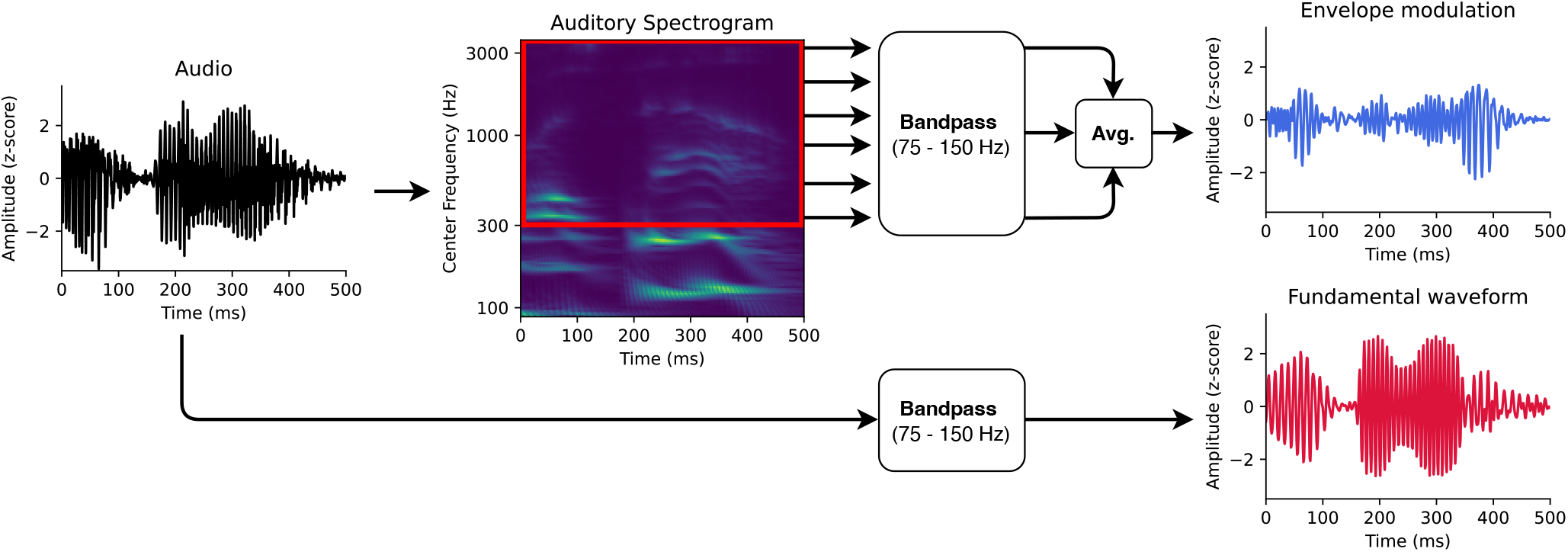
Auditory features for modelling the neural responses at the fundamental frequency. First, the fundamental waveform (red) was obtained by band-pass filtering the audio signal. Second, the high-frequency modulation of the envelope modulation was computed as well. The audio input was therefore transformed into an auditory spectrogram using a model of the auditory periphery. The frequency bins of the auditory spectrogram above 300 Hz were then filtered in the range of the fundamental frequency. The filtered bins were finally averaged to obtain the envelope modulation (blue).

The fundamental waveform is a waveform that oscillates at the fundamental frequency of the speaker’s voice. The original algorithm for computing the fundamental waveform was based on empirical mode decomposition (Huang and Pan (2006); Forte et al. (2017)). However, Etard et al. (2019) showed that direct band-pass filtering of the speech signal is considerably simpler, faster to compute and leads to the same result (Kulasingham et al. (2020); Van Canneyt et al. (2021b,a); Bachmann et al. (2021)). Here, we also employed a band-pass filter to extract the fundamental waveform from the voice recordings.

We used the software Praat (Boersma (2001)) and its Python interface Parselmouth (Jadoul et al. (2018)) to estimate the fundamental frequency in the voice recordings presented to the participants (44,100 Hz sampling rate). The mean fundamental frequency of the speaker was 107.2 Hz with a standard deviation of 24.8 Hz. The frequencies corresponding to the 5th and 95th percentiles of the speaker’s pitch distribution (75 Hz and 150 Hz, respectively) were used as corner frequencies of the bandpass filter. A FIR bandpass filter (7785th order, one-pass, zero-phase, non-causal, Hamming window, lower transition bandwidth: 18.7 Hz, upper transition bandwidth: 38.12 Hz) was then applied to filter the speech recordings. The resulting fundamental waveform was finally downsampled to 1 kHz to match the sampling rate of the EEG.

Since the neural response at the fundamental frequency might not emerge directly from the tracking of the speaker’s pitch but reflect the high-frequency envelope modulation, we used the latter as an additional feature in our analysis. The high-frequency envelope modulation was extracted from the audio signal as originally introduced (Kulasingham et al. (2020)). In particular, first, the audio signal was processed through a model of the auditory periphery reflecting the early stages of the auditory processing, including the cochlea, the auditory nerve and the subcortical nuclei (Chi et al. (2005))^3^ to obtain the auditory spectrogram with a millisecond temporal resolution (matching the sampling rate of the EEG).

The frequency bins of the obtained auditory spectrogram corresponding to the higher harmonics above 300 Hz were then band-pass filtered in the range of the fundamental frequency, between 75 Hz and 150 Hz (same FIR filter as for the fundamental waveform feature). The filtered signals were averaged to form the high-frequency envelope modulation feature. Similarly to the previous study employing the same pair of stimulus features (Kulasingham et al. (2020)), we found a negative correlation of *r* = −0.28 (Pearson’s) between the fundamental waveform and the high-frequency envelope modulation.

### 2.6 EEG modelling

Firstly, the acquired EEG data (1 kHz sampling rate) was band-pass filtered between 50 Hz and 280 Hz (265th order FIR one-pass, zero-phase, non-causal filter, Hamming window, lower transition bandwidth: 12.5 Hz, upper transition bandwidth: 70 Hz) and re-referenced to the average. The pre-processed EEG data and the stimulus features obtained from the corresponding speech signal were used to fit linear models following the methodology developed in Etard et al. (2019). EEG pre-processing and modelling pipelines were implemented through custom-written Python scripts using NumPy (Harris et al. (2020)), SciPy (Virtanen et al. (2020)) and MNE open-source packages (Gramfort et al. (2014)).

#### 2.6.1 Forward model

The forward models were designed to have complex coefficients. This approach allowed us to assess both the magnitude and the phase of the underlying neural response (Forte et al. (2017); Etard et al. (2019)). The complex forward model was designed to predict the multi-channel EEG response *r*(*t, c*) at channel *c* from the two stimulus features *f*_1_(*t*) and *f*_2_(*t*), where *f*_1_(*t*) represents the fundamental waveform and *f*_2_(*t*) the envelope modulation (Fig. 1). In particular, at each time instance *t*, the EEG signal was estimated as a linear combination of the stimulus features *f*_1_(*t*) and *f*_2_(*t*) as well as their Hilbert transforms 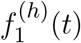 and 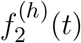 at a time lag *τ*:

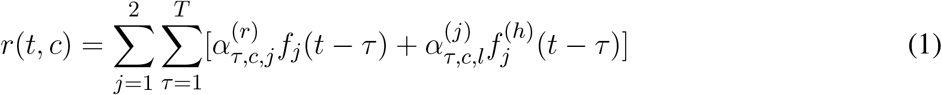

where 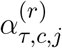 and 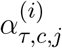 are real coefficients that can be interpreted as real and imaginary parts of a complex set of coefficients 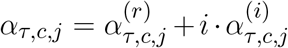. These coefficients are referred to as temporal response function (TRF) since they describe the time course of the neural response *r* to the two stimulus features *f*_1_ and *f*_2_. We note that the forward models is fitted using the two stimulus features simultaneously, analogously to Kulasingham et al. (2020).

We used *T* = 750 time lags ranging from −250 ms (i.e. the stimulus is preceded by the EEG signal, thus anticausal) up to 499 ms. We chose a broad range of time lags, including a latency range typical for cortical responses, to include both early and putative late responses. The model coefficients were obtained using ridge regression (Hastie et al. (2009)) with a regularization parameter λ = *λ_n_* · *e_m_*, where λ*_n_* is a normalized regularization parameter and *e_m_* is the mean eigenvalue of the covariance matrix, to which the regularization was added (Biesmans et al. (2017)). For the forward model, we used a fixed normalized regularization parameter of λ*_n_* = 1. Prior to fitting the model, each EEG channel and the stimulus features were standardized by subtracting their mean and dividing them by their standard deviation.

A complex forward model was computed separately for each participant. The subject-specific models were then averaged to obtain a population-averaged model. The magnitudes of the complex coefficients were computed by taking their absolute values, and the phases by computing their angles. To summarize the contribution of different time lags, the magnitudes of the population-averaged model were additionally averaged across channels to obtain a single value per time lag. This value, reflecting the contribution of each time lag to the model, allowed us to estimate the latency of the predominant neural response.

To assess the significance of the forward model, we established null models using time-reversed stimulus features. Due to the mismatch between the speech features and the EEG signal, the null models were purposefully designed to reflect no meaningful brain response across the entire range of time lags. One null model was obtained for each subject. We bootstrapped the population-level null models by re-sampling null models across participants (with replacement), averaging them and computing their magnitudes across time lags in the same way as for the actual forward model. This procedure was repeated 10,000 times to form a distribution of null model magnitudes across time-lags. We therefrom computed an empirical *p*-value for each time lag by counting how many values from the null distribution exceeded the actual forward model for each time lag. Finally, the obtained *p*-values were corrected for multiple comparisons using the Benjamini-Yekutieli method (Benjamini and Yekutieli (2001)).

#### 2.6.2 Backward model

Backward models were designed to reconstruct the two stimulus features *f*_1_ (*t*) and *f*_2_(*t*) from the time-lagged multi-channel EEG response *r*(*t, c*). In particular, for each time instance *t*, the stimulus features were reconstructed as follows:

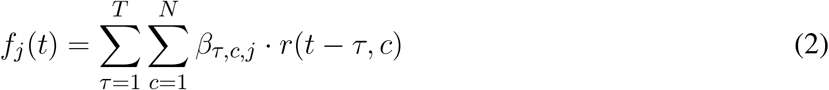

where *β_τ,c,j_* are real-valued model coefficients, *j* ∈ {1, 2} denotes the stimulus feature, *c* represents the index of the EEG channel, and *τ* is a time lag between the auditory stimulus features and the EEG recording. Here, we used *T* = 55 time lags ranging from −5 ms (i.e. the EEG signal preceded the stimulus) to 49 ms. We only used real and not complex model coefficients, since the use of the former did not impact the reconstruction performance but greatly decreased the computational cost. The coefficients of the backward models were obtained in the same way as for the forward model using ridge regression. We evaluated 51 logarithmically-spaced normalized regularization parameters λ*_n_* ranging from 10^−10^ to 10^10^. Analogously to the forward models, the backward models were fitted using two stimulus features simultaneously.

Backward models were evaluated through five-fold cross-validation (Hastie et al. (2009)). In particular, all the available data were split into five folds of the same duration of approximately eight minutes. Four folds were used to train the backward model, and the remaining one was kept aside for evaluating the model. Each time, 51 models, one for each regularization parameter, were trained and evaluated.

To investigate how the amount of available data influenced the reconstruction performance, we split the testing data into either segments of arbitrary lengths or according to the word boundaries. The performance of each model was quantified by computing Pearson’s correlation coefficient between the reconstructed stimulus features and the actual one. After evaluating the models on all segments, another fold of the data was selected as the testing set. The procedure was repeated five times until all the available data was used. This yielded reconstruction scores that reflected the strength of the neural response.

To test whether segmenting the evaluation data according to word boundaries yielded different performances in reconstructing the fundamental waveform, we compared it to the reconstruction scores obtained using segments of arbitrary duration agnostic of word onsets. For the latter, we considered six different durations of the arbitrary testing segments. Since the averaged word duration was 260 ms, we chose the fixed evaluation segment durations to be 100 ms, 260 ms (the mean word duration), 310 ms (the median word duration), 1 s, 10 s and 30 s. We evaluated the backward models as specified above for all 13 subjects. In particular, reconstruction scores from all testing segments across all the folds were averaged to summarize the model performance for each subject. For this analysis, we used the fixed normalized regularization parameter λ*_n_* = 1.

We thereby obtained 13 averaged reconstruction scores (one per subject) for each stimulus feature (fundamental waveform and envelope modulation) and for each segment duration. For each stimulus feature, we performed the Friedman test, a non-parametric equivalent of ANOVA, to assess whether at least one of the evaluation segment lengths yielded different reconstruction scores from the others. Then, we performed a post-hoc test on the results for each pair of segment durations through the Wilcoxon signed-rank test. In addition, the reconstruction scores for the two stimulus features were compared for each segment duration. The *p*-values obtained from the above tests were corrected for multiple comparisons using the Benjamini-Yekutieli method (Benjamini and Yekutieli (2001)).

The null models that represented the chance-level reconstruction scores were obtained in the same way as described above, but using the time-reversed stimulus features. Following the same reasoning as for the forward model, these models contained no actual brain response and estimated the chance-level reconstruction scores.

### 2.7 Word-level features

We used seven distinct word-level features to study the neural response at the fundamental frequency of the continuous narratives (Fig. 2A). Four linguistic features were developed in Weissbart et al. (2020) and are openly available on figshare.com^4^. In short, the transcriptions of the stories presented to the participants were processed through a language model to obtain the frequency, surprisal and precision of each word. The word frequency reflects the probability of a word out of context and was computed from Google N-grams by taking only the unigram values. As a result, this feature estimated the unconditional probability of the occurrence of a word *P*(*w*). To match the remaining information-theoretic features, we computed the negative logarithm of this probability *-ln*(*P*(*w*)), and refer to this feature as the ‘inverted word frequency’ in the following. Importantly, less frequent words were therefore assigned a higher inverted word frequency, and more frequent words were assigned lower values.

**Figure 2.**
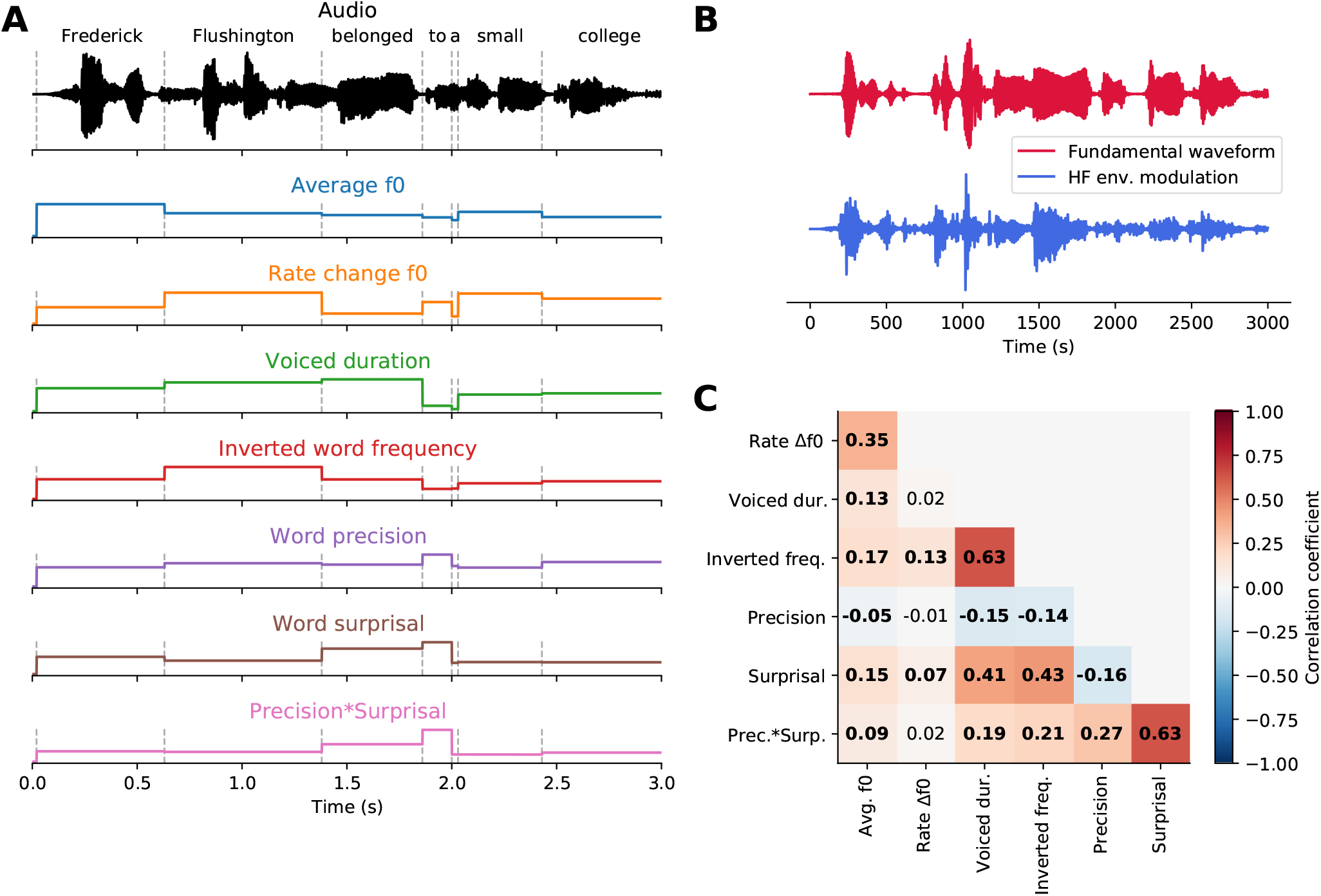
Word-level features. **(A)**, An exemplary part of a speech signal (black). Dashed vertical lines represent word onsets. Seven features (coloured) were used to describe each word in the stories presented to the participants. Three of them – the averaged fundamental frequency (f0), the rate of f0 change and the duration of the voiced part – were acoustic features based on the voiced parts of each word. The remaining four features – inverted word frequency, word surprisal, word precision and the interaction of precision and surprisal – were derived from a language model and characterized linguistic properties of each word. **(B)**, The two stimulus features extracted from the exemplary audio segment, the fundamental waveform (red) and the high-frequency (HF) envelope modulation (blue). **(C)**, Pairwise correlations between wordlevel features. Significant correlations (*p* < 0.05, after correcting for multiple comparisons using the Benjamini-Yekutieli method) are denoted in bold.

In contrast to the inverted word frequency, the word precision and surprisal were derived from conditional probabilities of a particular word given the preceding words. In particular, the probability of the *n*th word *w_n_* can be expressed as *P*(*w_n_*|*w*_1_, *w*_2_, …, *w*_*n*–1_). Word surprisal quantifies the information gain that an upcoming word generates given the previous words and reflects how unexpected the word is in its context. Here, the surprisal of the word *w_n_* was computed as the negative logarithm of its conditional probability:

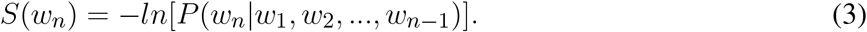

In contrast to the word surprisal, the word precision reflects the confidence about the prediction of the next word given the previous words. Here, the word precision was computed as the inverse word entropy, [*E*(*w_n_*)]^−1^. On its own, the word entropy represents the uncertainty of predicting the next word *w_n_* from the past context (*w*_1_, *w*_2_,…, *w*_*n*–1_), and is formulated as:

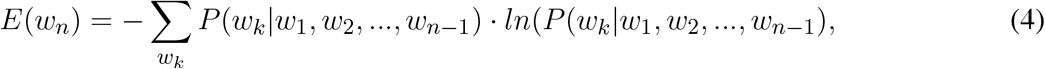

where *w_k_* denotes the *k*th word from the text corpus.

Finally, to investigate a possible modulating effect that precision may have on surprisal, an interaction term was obtained by multiplying precision with surprisal. This feature can be interpreted as a confidence-weighted surprisal or a surprisal-dependent precision.

The conditional probabilities, required for computing the word surprisal and precision, were obtained from a recurrent neural network (RNN) language model introduced in Mikolov et al. (2011). The model was designed to predict the current word *w_n_* given the previous words *w*_1_, *w*_2_,…, *w*_*n*–1_. Firstly, embeddings of words in the input text were obtained using pre-trained global vectors for word representation (GLOVE) trained on the Wikipedia 2014 and Gigaword 5 datasets (Pennington et al. (2014)). The obtained embeddings were projected to 350 recurrent units forming the hidden layer of the model. The output layer of the model was a softmax function, from which the word probabilities were computed. Such model was trained on the *text8* dataset, consisting of 100 MB of text data from Wikipedia (Mahoney (2011)), using backpropagation through time and a 0.1 learning rate. Prior to the training, the text data was cleaned to remove punctuation, HTML, capitalization and numbers. In addition, to facilitate model training, the vocabulary was limited to the 35,000 most common words in the dataset. The remaining rare words were assigned an *‘unknown*’ token. For more implementation details of the model itself and its training, please see Weissbart et al. (2020) where the method was originally developed.

Having obtained the above-described linguistic features, each word in the story was aligned to the acoustic signal using a forced alignment algorithm implemented in the Prosodylab-Aligner software (Gorman et al. (2011)). Subsequently, we computed three additional acoustic features for each word. In particular, we used the Praat & Parselmouth Python interface (Boersma (2001); Jadoul et al. (2018)) to obtain the evolution of the speaker’s fundamental frequency across the story recording.

For each word, we then computed the duration of its voiced part, its mean fundamental frequency and the rate of the change in the fundamental frequency. The latter feature was obtained by averaging the absolute value of the first derivative of the fundamental frequency’s time course across the voiced duration,as described by Van Canneyt et al. (2021b). Including these three features in our analysis allowed us to control for purely acoustic modulation of the neural response at the fundamental frequency.

### 2.8 Stepwise hierarchical regression

We first determined the strength of the neural response at the fundamental frequency for the ith word. To this end, the backward models for each participant were evaluated to obtain a reconstruction score for each word in the story (*N* = 6, 345). 100 words did not contain a voiced part, and were therefore discarded from further analysis. For each remaining word, we obtained 51 reconstruction scores, one for each normalized regularization parameter λ*_n_*. We picked the optimal regularization parameter λ*_model_*, leading to the best reconstruction.

To control for overfitting, the same procedure was applied to the backward null models that did not contain a meaningful brain response. Possible inflation of the reconstruction score *r*(*i*) from overfitting was corrected by subtracting the score obtained by the null model *r_null_*(*i*) from that of the actual decoder *r_model_*(*i*):

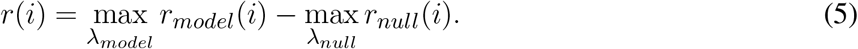

The above procedure was applied independently to the reconstruction scores obtained for the two stimulus features, the fundamental waveform and the envelope modulation.

We note that the optimal word-level regularization parameter was picked independently for the actual (*λ_model_*) and the null model (*λ_null_*). Controlling for overfitting in this empirical manner allowed to avoid pre-selecting a fixed regularization parameter, which could either inflate or deflate reconstruction scores. However, as an additional control, we also computed the reconstruction scores with a pre-selected regularization parameter of λ*_model_* = 1. The resulting reconstruction scores were not significantly different from those obtained with the procedure outlined above (*p* > 0.382, Wilcoxon signed-rank test). Having computed the single-word reconstruction scores for each participant, we averaged them across the subjects for each word in the story to obtain population-level single-word reconstruction scores.

We then investigated whether the word-level acoustic and linguistic features modulated the early neural response at the fundamental frequency, that is, whether they modulated the single-word reconstruction scores. To this end, we first standardized both the single-word reconstruction scores *r* and the word-level features *x* by subtracting their mean and dividing them by their standard deviation. In addition, we used the isolation forest (Liu et al. (2008)), an unsupervised algorithm based on the random forest, for detecting outliers and anomalies.

In this method, data points corresponding to the words in the stories and described by the word-level features and the reconstruction scores (eight descriptors) were processed through a set of 1,000 random trees established based on the dataset statistics. The algorithm measured the average path it took each data point to traverse from the roots of the trees in the random forest to their leaf nodes. Since the outliers contained extreme descriptor values, they reached leaf nodes earlier, yielding shorter paths. The threshold for the path length qualifying the data point as an outlier was determined automatically based on the dataset-wide statistics. Here, we used the implementation of the method included in the scikit-learn Python package (Pedregosa et al. (2011)). Following the outlier removal, 5,732 data points corresponding to different words in the stories remained.

We then related the single-word reconstruction scores to the word-level features through stepwise hierarchical regression. The approach was inspired by the stepwise and hierarchical regression commonly used for feature selection in multiple regression models (Lewis (2007)). However, neither of the standard approaches is suited for the cases in which the explanatory variables exhibit a degree of multicollinearity. Since the word-level linguistic and acoustic features were indeed correlated (Fig. 2C), we employed a stepwise approach based on the expected effect size for each feature, in a hierarchical manner.

The *n* word-level features were first ordered from that with the highest expected predictive power, *x*^(1)^, to that with the lowest expected predictive power, *x*^(*n*)^. In the first step of the procedure, the word-level feature *x*^(1)^ from the ordered list was used to fit a linear model to predict word-level reconstruction scores 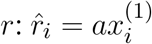 with a coefficient *a*, assuming that *r* and *x* were standardized. In this equation, *x_i_* denotes the world-level features of the *i*th word, and *r_i_* its reconstruction score.

The feature was then projected out from the word-level reconstruction scores *r* by subtracting the estimated word-level reconstruction scores 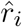 from the actual ones: 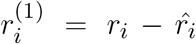. The residual reconstruction scores 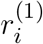 were used as a response variable for fitting the next linear model using the next word-level feature *x*^(2)^ from the ordered list. The process was repeated until all available word-level features were used.

By projecting out the predictions obtained from subsequent word-level features, we assured that a possible predictive contribution of a feature with a lower expected predictive power did not result from that of a feature with a higher expected predictive power due to shared variance. The stepwise hierarchical regression therefore constituted a conservative manner to ensure that any contributions from features with lower expected predictive power were indeed real, and that their significance was not inflated due to the shared variance with features with a higher expected predictive power.

To further reduce the influence of the extreme data points on the model coefficients, we fitted the linear models in the stepwise hierarchical regression through robust regression, using Huber weighting of the residuals (Andrews (1974)), instead of the ordinary least squares regression.

Regarding the ordering of the different features, due to the reported significant impact of the acoustic features on the neural response at the fundamental frequency (Saiz-Alía et al. (2019); Saiz-Alía and Reichenbach (2020); Van Canneyt et al. (2021b)), we prioritized these above the linguistic features that likely have a weaker impact. We adopted the following ordered list of the word-level features for the stepwise hierarchical regression: (1) average fundamental frequency f0, (2) rate of the change in f0, (3) duration of the voiced part, (4) inverted word frequency, (5) word precision, (6) word surprisal, and (7) precision x surprisal.

The stepwise hierarchical regression was applied for both stimulus features, the fundamental waveform and the envelope modulation, for which the reconstruction scores were computed. Each time, the output of the procedure was a set of seven linear models corresponding to the seven word-level features. The p values reflecting the significance of each model coefficient were corrected for multiple comparisons using the Benjamini-Yekutieli method (Benjamini and Yekutieli (2001)).

The methods described above were implemented via custom-written Python scripts using NumPy (Harris et al. (2020)), SciPy (Virtanen et al. (2020)) and statsmodels open-source packages (Seabold and Perktold (2010)).

## 3 RESULTS

### 3.1 Relations between the word-level acoustic and linguistic features

We computed Pearson’s correlation coefficient between each pair of features (Fig. 2C). The obtained correlation ranged from −0.157 to 0.632. The highest correlation coefficient (*r* = 0.632) emerged between surprisal and the interaction of surprisal and precision. Another significant positive correlation (*r* = 0.431) arose between inverted word frequency and surprisal, indicating that less frequent words tended to be more surprising. Similarly, a positive correlation (*r* = 0.406) between voiced duration and surprisal showed that more surprising words had longer voiced parts and were presumably longer overall. A positive correlation (*r* = 0.632) between inverted word frequency and voiced duration indicated that less frequent words tend to be longer. The remaining correlations between features were comparatively small, between −0.157 and 0.274. The rate of fundamental frequency change was the least correlated with other features. It was only significantly correlated with the inverted word frequency (*r* = 0.129).

### 3.2 Early neural response at the fundamental frequency

We investigated the neural response at the fundamental frequency through the temporal response functions (TRFs) obtained from the forward model (Haufe et al. (2014)). In particular, we examined the model coefficients associated with the two considered stimulus features, the fundamental waveform and the high-frequency envelope modulation.

For the neural response to the fundamental waveform (Fig. 3 A,B,C), the channel-averaged TRFs yielded significant responses for short delays between 9 ms - 12 ms, with a peak at 11 ms. The TRFs at the peak delay showed the highest magnitudes in the central-frontal and occipital regions, as well as at the mastoid electrodes. The phase relationship of the model coefficients at the delay of 11 ms exhibited a phase shift of approximately π between the frontal and occipital areas, and a slightly larger phase difference between the central-frontal and the mastoid electrodes.

**Figure 3.**
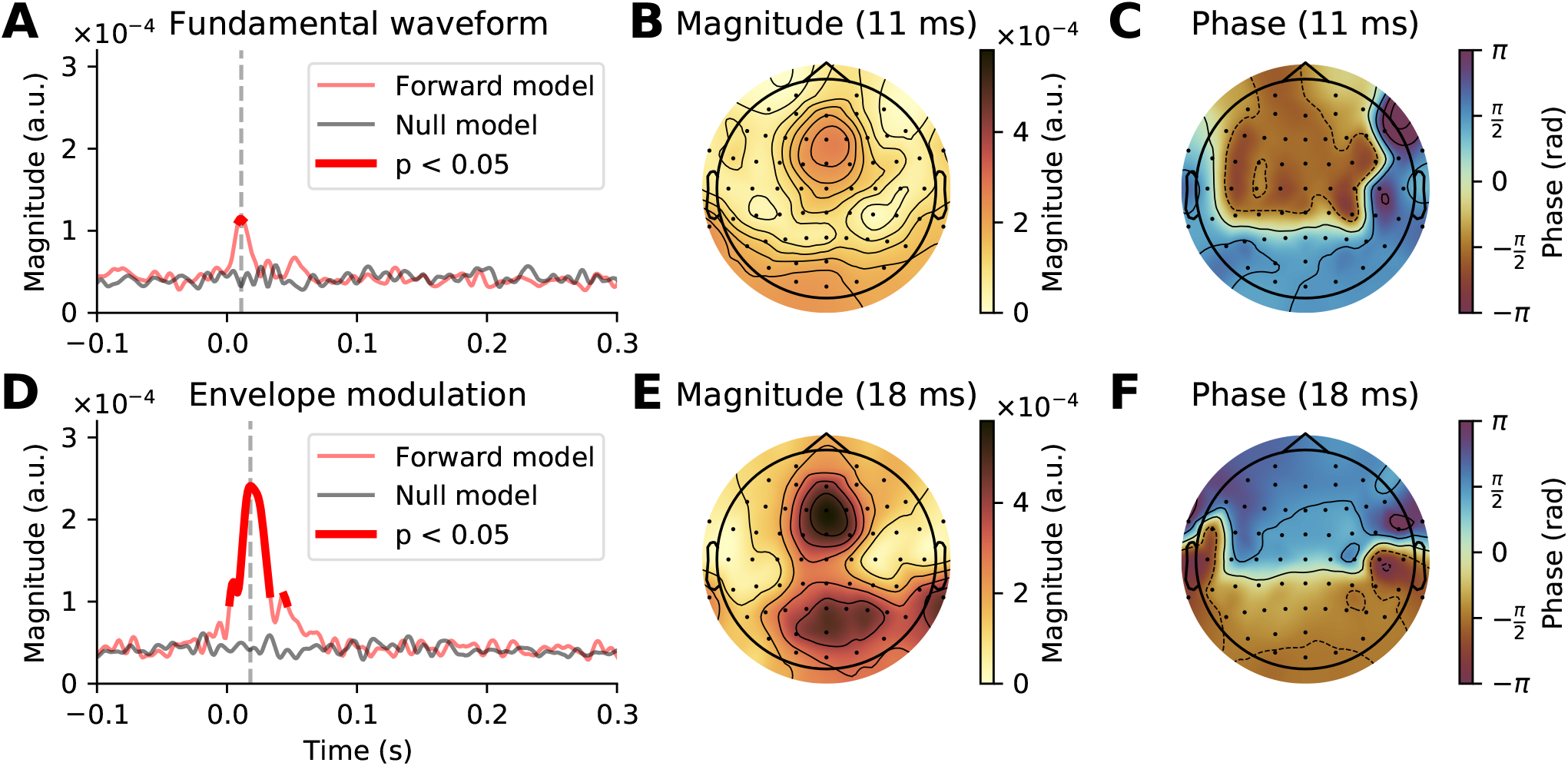
Early neural response at the fundamental frequency of continuous speech. **(A - C)**, Response to the fundamental waveform. **(A)**, The magnitude of the complex coefficients of the forward model (red), averaged across EEG channels and subjects exhibited a peak at an early latency of 11 ms (grey dashed line). The comparison of the complex TRF magnitudes to a null model (black solid line) showed that significant responses emerge only at latencies around the peak latency, between 9 ms and 12 ms (thicker red line, *p* < 0.05, corrected for multiple comparisons). **(B)**, At the peak latency of 11 ms, the largest contribution to the TRF came from central-frontal and occipital areas, as well as from the mastoid electrodes. **(C)**, The phase of the model coefficients indicated a phase shift of approximately *π* between the frontal area on the one hand and the occipital and mastoid electrodes on the other hand. **(D - F)**, Neural response to the high-frequency envelope modulation. **(D)**, The average magnitude of the complex TRF coefficients was substantially larger than that of the response to the fundamental waveform. In particular, the coefficients of the model significantly exceeded the chance level between 2 ms to 33 ms and 44 ms to 46 ms, with the peak magnitude at 21 ms. **(E, F)**, At the peak latency of 18 ms the TRFs exhibited similar topographic patterns to those obtained for the response to the fundamental waveform.

For the response to the envelope modulation (Fig. 3 C,D,E), the channel-averaged TRFs showed significant contributions between 2 ms - 33 ms as well as 44 ms - 46 ms, with a peak at 18 ms. Notably, the averaged magnitudes of the models coefficients for this response were over two times larger than for the response to the fundamental waveform. Despite the peak magnitude occurring later, the topographical pattern of the model coefficients, both in terms of magnitudes and phases, were similar to that obtained for the response to the fundamental waveform. In particular, the largest magnitudes were obtained for frontal and occipital regions, with a phase difference of approximately π between them.

### 3.3 Reconstruction of the stimulus features from EEG

We then assessed the reconstruction of the stimulus features from the EEG recordings using backward models (Fig.4). In particular, we investigated whether the reconstruction performance varied with the duration of the speech segment on which the models were tested.

**Figure 4.**
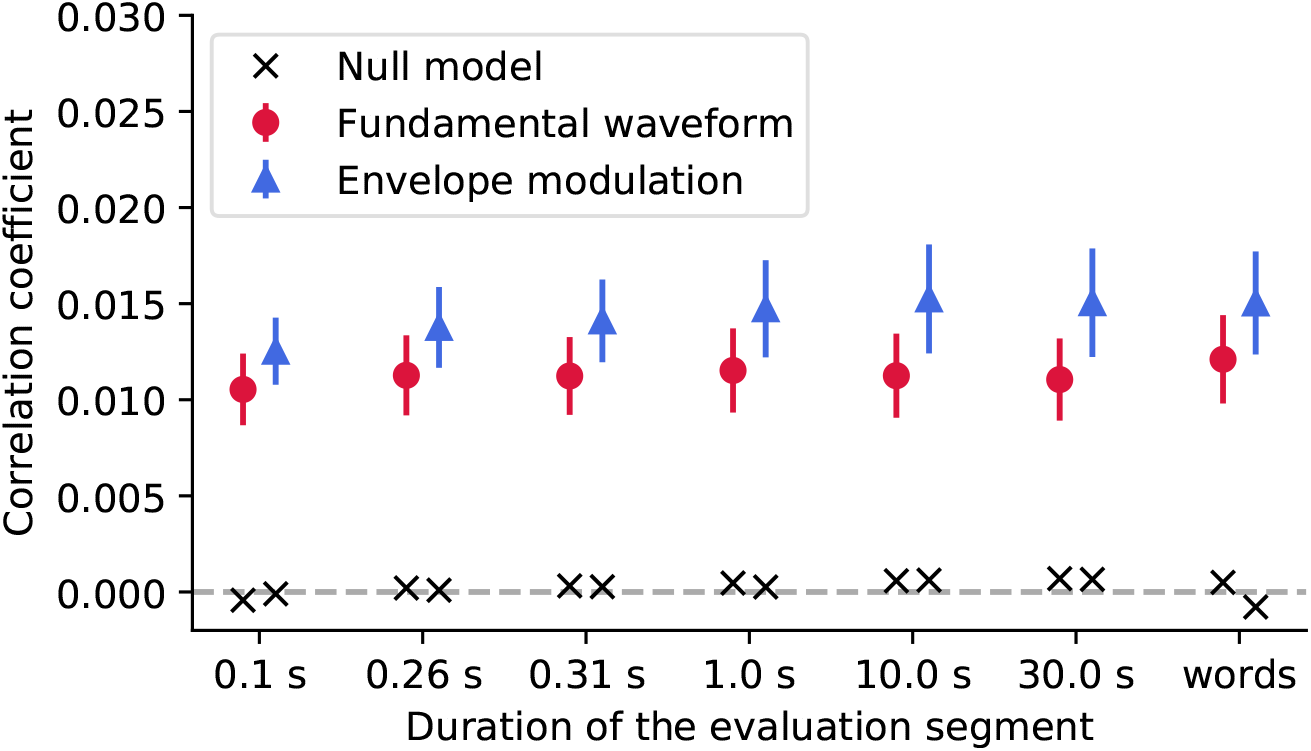
We evaluated the reconstruction of the stimulus features from the EEG recordings using segments of different duration, including segments aligned with the word boundaries (*words*). For each segment duration, the population-averaged reconstruction scores obtained for the fundamental waveform feature are denoted with red circles, and for the envelope modulation with blue triangles. The error bars correspond to the standard error of the mean across participants. For both features, the reconstruction scores yielded much higher correlation coefficients as compared to their respective null models (black crosses).

The Friedman test was applied to the reconstruction scores to assess whether either of the segment duration yields significantly better reconstructions than the other. For neither of the features, the test yielded significant results (*p* > 0.101, corrected for multiple comparisons), indicating that none of the considered segment durations, for neither of the two features, yielded different reconstruction scores from the rest.

However, the reconstruction scores obtained for the envelope modulation feature were overall higher than those obtained for the fundamental waveform. In particular, the differences were significant for some segment durations (Wilcoxon signed-rank test, 0.1 s - *p* = 0.055; 0.26 s - *p* = 0.037; 0.31 s - *p* = 0.037; 1.0 s - *p* = 0.037; 10.0 s - *p* = 0.052; 30.0 s - *p* = 0.049; *words* - *p* = 0.037, all corrected for multiple comparisons). Overall, the slightly higher reconstruction score for the envelope modulation feature is in agreement with the forward model, whereby the envelope modulation features also yielded higher neural response (reflected by the forward model coefficients).

### 3.4 Modulation of the early neural response at the fundamental frequency through acoustic and linguistic features

We used stepwise hierarchical regression to investigate the acoustic and linguistic modulation of the early neural response at the fundamental frequency of continuous speech. Through this method, we predicted the word-level reconstruction scores of the backward model, reflecting the strength of the neural response, from the seven word-level features (Fig. 2A).

We first predicted the reconstruction scores related to the fundamental waveform (Table 1, Fig. 5). Of the considered acoustic word-level features, the average fundamental frequency (f0) of a word’s voiced part (−0.075, *p* = 2 · 10^−7^) and the rate of change of the fundamental frequency (−0.083, *p* = 2 · 10^−5^) both yielded significant model coefficients with similar values. The negative values of the model coefficients showed that higher average fundamental frequency and higher associated variability leads to less neural tracking of the fundamental waveform. However, neither the duration of a word’s voiced part nor any of the four considered linguistic features had a significant influence on the reconstruction scores (*p* > 0.3).

**Table 1.** Word-level modulation of the neural response to the fundamental waveform. The table presents the model coefficients obtained from stepwise hierarchical regression, using the reconstruction scores of the fundamental waveform. It details the model coefficient (Coeff.), the standard error (SE), the 95% confidence interval (CI), the *z* statistic (*z*) and the *p*-value after the FDR correction for multiple comparison using the Benjamini-Yekutieli method. Word-level features that yield a significant contribution are highlighted in bold, as well as through an asterisk.

| Feature | Coeff. | SE | 95% CI | $z$ | $p$ (FDR) |
| --- | --- | --- | --- | --- | --- |
| **Average f0 (\*)** | -0.075 | 0.013 | (-0.100; -0.049) | -5.716 | $2 \cdot 10^{-7}$ |
| **Rate change f0 (\*)** | -0.083 | 0.018 | (-0.117; -0.049) | -4.728 | $2 \cdot 10^{-5}$ |
| Voiced duration (n.s.) | -0.013 | 0.013 | (-0.039; 0.013) | -0.963 | 1.000 |
| Inverted word frequency (n.s.) | -0.027 | 0.014 | (-0.054; -0.001) | -1.964 | 0.300 |
| Word precision (n.s.) | -0.046 | 0.038 | (-0.122; 0.029) | -1.205 | 0.829 |
| Word surprisal (n.s.) | -0.022 | 0.014 | (-0.047; 0.009) | -1.540 | 0.561 |
| Precision x Surprisal (n.s.) | -0.003 | 0.023 | (-0.047; 0.047) | -0.125 | 1.000 |

**Figure 5.**
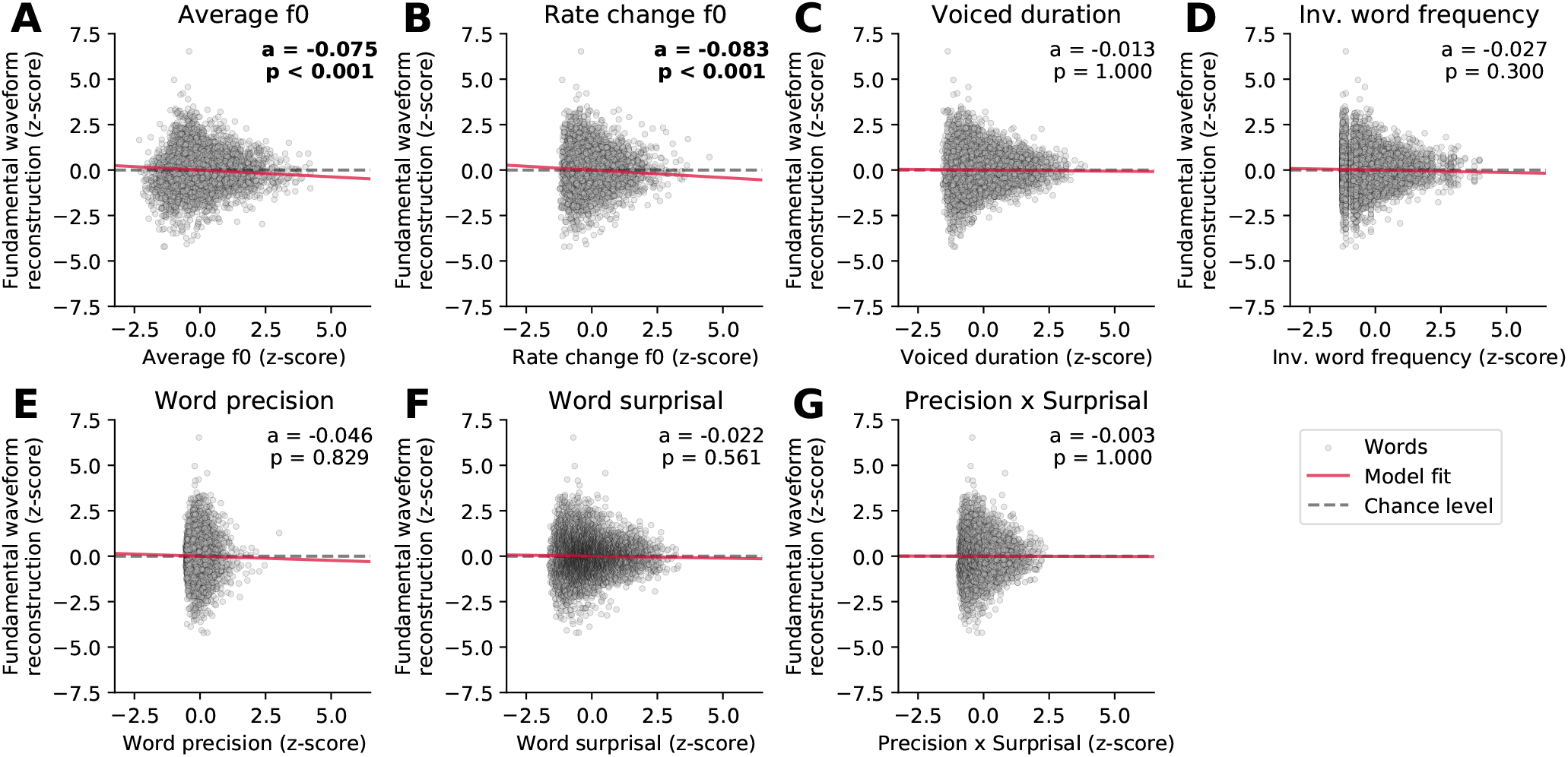
Dependency of the strength of the neural response to the fundamental waveform on the different word-level features. Panels (**A** - **G**) show the standardized single-word reconstruction scores averaged across 13 participants against the seven standardized word-level features. Each scatter plot shows data points corresponding to 5,732 words from the story presented to the participants during the EEG acquisition. The slopes of the red lines correspond to the coefficients obtained from the stepwise hierarchical regression. Each panel also depicts the model coefficient (a) for a given feature and the associated *p*-value (FDR-corrected). The gray dashed lines are horizontal and indicate no dependency.

We then investigated which word features could predict the reconstruction scores of the high-frequency envelope modulation (Table 2, Fig. 6). As for the neural response to the fundamental waveform, both the average fundamental frequency and its rate of change significantly modulated the reconstruction scores. In particular, the average fundamental frequency was related to an even larger negative coefficient (−0.170, *p* =1 · 10^−37^) than for the neural response to the fundamental waveform. In contrast, the rate of change of the fundamental frequency within words led to a slightly smaller negative coefficient (−0.070, *p* = 7· 10^−4^). The duration of the voiced portion of each word did not significantly modulate the reconstruction scores.

**Table 2.** Word-level modulation of the neural response to the high-frequency envelope modulation. The table presents model coefficient obtained from stepwise hierarchical regression. It lists the model coefficient (Coeff.), the standard error (SE), the 95% confidence interval (CI), the *z* statistic (*z*) and the *p*-value after the FDR correction for multiple comparison using the Benjamini-Yekutieli method for the different word-level features. Word-level features that yield significant contributions (*p* < 0.05) are highlighted in bold, as well as through an asterisk.

| Feature | Coeff. | SE | 95% CI | $z$ | p (FDR) |
| --- | --- | --- | --- | --- | --- |
| **Average f0 (\*)** | -0.170 | 0.013 | (-0.195; -0.144) | -13.048 | $1 \cdot 10^{-37}$ |
| **Rate change f0 (\*)** | -0.070 | 0.018 | (-0.105; -0.036) | -3.967 | $7 \cdot 10^{-4}$ |
| Voiced duration (n.s.) | -0.015 | 0.013 | (-0.041; 0.010) | -1.169 | 0.880 |
| **Inverted word frequency (\*)** | -0.040 | 0.014 | (-0.067; -0.013) | -2.889 | 0.023 |
| Word precision (n.s.) | -0.054 | 0.039 | (-0.129; 0.022) | -1.388 | 0.749 |
| Word surprisal (n.s.) | -0.007 | 0.014 | (-0.035; 0.020) | -0.497 | 1.000 |
| Precision x Surprisal (n.s.) | 0.004 | 0.023 | (-0.041; 0.049) | 0.179 | 1.000 |

**Figure 6:**
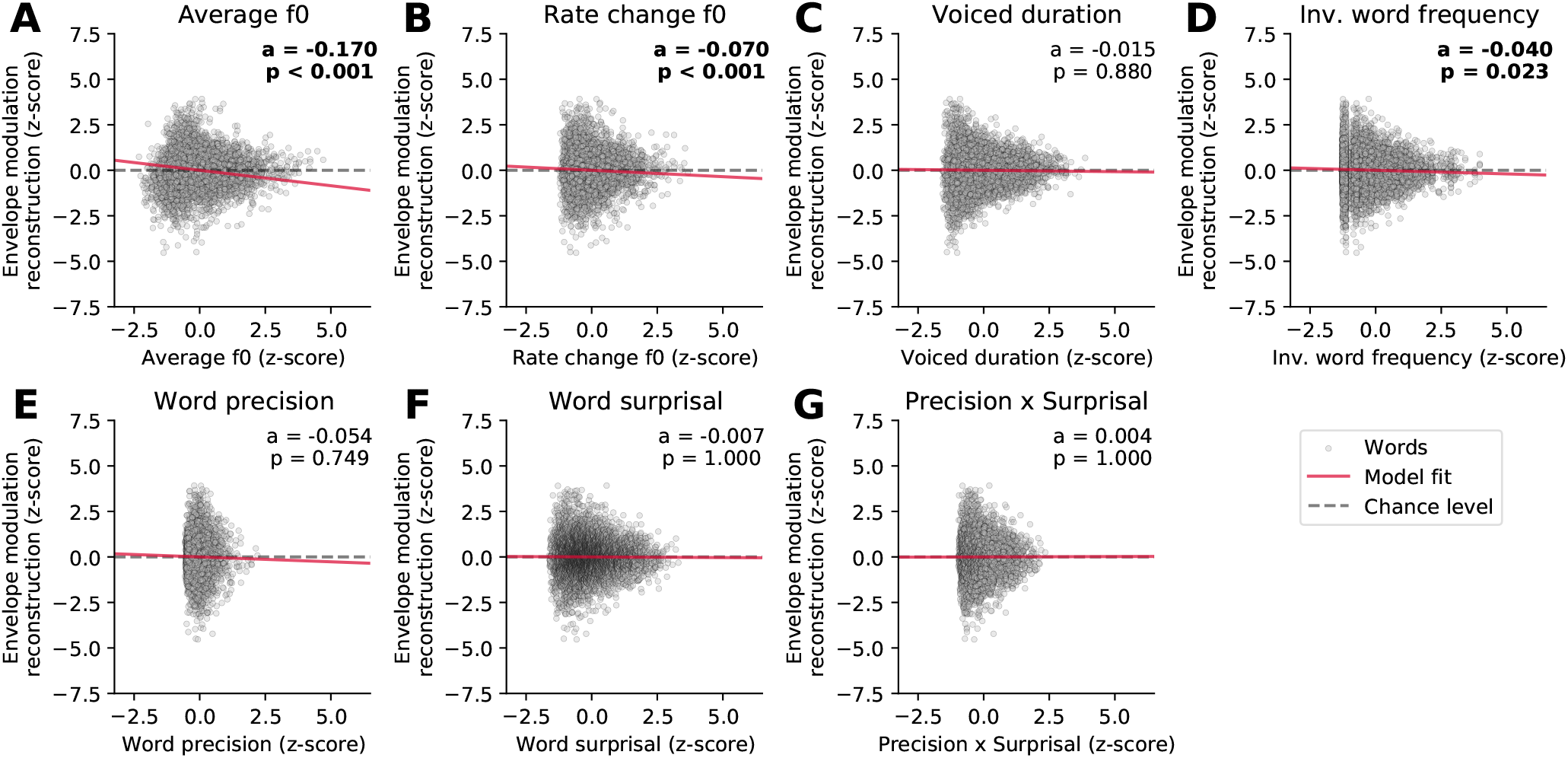
Dependency of the strength of the neural response to the high-frequency envelope modulation on the different word-level features. The world-level features and the reconstruction scores were standardized. The data points in each plot correspond to 5,732 words, and the slopes of the red lines show the coefficients of the stepwise hierarchical regression. We also detail the model coefficient (a) and the associated *p*-value (FDR-corrected). The gray dashed lines are horizontal and indicate no dependency.

Importantly, the inverted word frequency, the 4*^th^* feature in the hierarchy, was a significant predictor of the reconstruction scores related to the envelope modulation. This linguistic feature was assigned a small, but significant, negative coefficient (−0.040, *p* = 0.023), indicating that less frequent words (with higher inverted frequency value) led to less neural tracking of the high-frequency envelope modulation. In contrast, none of the context-dependent word-level features (precision, surprisal and their interaction) yielded significant model coefficients (*p* > 0.749).

## 4 DISCUSSION

We showed that the word-level early neural response at the fundamental frequency of natural speech is modulated predominantly by acoustic features, but also by one of the four considered linguistic features, the inverted word frequency. Previous studies have shown significant modulation of the neural response at the fundamental frequency by acoustic differences between different speakers (Saiz-Alía et al. (2019); Saiz-Alía and Reichenbach (2020); Van Canneyt et al. (2021b)). Here, we extended these findings by showing that the same effect persists for fluctuation of acoustic properties between distinct words produced by the same speaker.

The fundamental frequency of the speech that we employed varied between 75 Hz and 150 Hz. The neural response occurred accordingly at comparatively high frequencies. It could be evoked either directly by the fundamental frequency or by the high-frequency modulations of higher harmonics (Kulasingham et al. (2020)). We considered both of these features and included them in our EEG modelling framework. We found that the response associated with the high-frequency envelope modulation was considerably stronger than that associated with the fundamental waveform, as observed previously in MEG recordings (Kulasingham et al. (2020)). We furthermore found that the response associated with the fundamental waveform occurred earlier, around 11 ms, as compared to that associated with the high-frequency envelope modulation, at about 18 ms and at 44 ms.

The neural response at the fundamental frequency, as well as the related FFR to pure tones, is mostly attributed to the subcortical nuclei, the inferior colliculus and the medial geniculate body (Chandrasekaran and Kraus (2010); Skoe and Kraus (2010)). However, recent MEG and EEG investigations have also identified a cortical contribution, in particular at frequencies below 100 Hz (Coffey et al. (2016, 2017); Bidelman (2018); Gorina-Careta et al. (2021)). Regarding the measurements presented here, the earlier response associated to the fundamental waveform (at a delay of 11 ms) may have resulted predominantly from the brainstem and midbrain, as suggested by the low latency, high frequency and sensor-space topography of the response.

The later response to the high-frequency envelope modulation (at a delay of about 18 ms as well as at 44 ms) might, however, represent cortical contributions. Previous MEG recordings did indeed find cortical responses to the high-frequency envelope modulation of speech at a delay of about 40 ms (Kulasingham et al. (2020)). The latencies of this response are similar to those in the auditory middle latency response that is assumed to originate in Heschl’s gyrus (Borgmann et al. (2001); Liegeois-Chauvel et al. (1994); Yoshiura et al. (1995)). However, due to the considerable autocorrelation of the stimulus features, our measurements did not allow us to further resolve these different neural components in the temporal domain, and the relatively low spatial resolution of our EEG measurements prevented us from more detailed spatial source localization as well. We could therefore not distinguish whether the modulation of the neural response at the fundamental frequency through the acoustic and linguistic features occurred at the subcortical level, at the cortical level, or at both.

Irrespective of the precise neural origin of the response, however, the small latency of the response implies that its modulation through the linguistic features likely results from feedback from higher cortical areas at which the linguistic information in speech is processed. If the relevant contribution to the neural response originates from subcortical areas, such as the inferior colliculus, this would require corticofugal feedback to be involved in linguistic processing. If a cortical source of the neural response was modulated by the linguistic features, then the linguistic processing would involve feedback projections between different cortical areas.

To investigate the modulation of this neural response by the different acoustic and linguistic word-level features, we developed the methodology to estimate the neural response at the fundamental frequency at the word level. We tested the validity of our method by comparing the accuracy of the stimulus feature reconstruction by the backward models for different lengths of audio segments. As expected, since the models were optimized on the same training data, the segmentation of the evaluation set did not impact the feature reconstruction scores. The reconstruction performance for the envelope modulation feature was however systematically slightly higher for the envelope modulation feature for all considered durations. This matches the forward modelling results whereby the neural response, reflected by the model coefficients, was much weaker for the fundamental waveform feature.

We employed three acoustic features, the average fundamental frequency, the rate of the fundamental frequent change and the duration of the voiced portion of a word. As discussed above, the first two word-level acoustic features strongly modulated the neural response at the fundamental frequency, both that related to the fundamental frequency and that related to the high-frequency envelope modulation. The stronger modulation of the neural tracking of the high-frequency envelope modulation might be explained by the stronger neural response to this feature.

The duration of the voiced portion of words in the story, however, had no significant impact on the neural tracking of either of the stimulus features. We note that we excluded entirely voiceless words from the analysis, since we could not infer a neural response for those. The neural response at the fundamental frequency is accordingly relatively similar for shorter and for longer voiced durations. Although longer voice durations will allow a better estimate of the neural response, that is, at a better signal-to-noise ratio, the response itself is indeed expected to stay constant. In other words, while longer segments of training data will lead to a more accurate backward model, the model’s inference capability is independent of the duration of the data on which it is tested. This result concurs with our finding that the strength of the neural response remains unaffected by the duration of the data on which it is evaluated (Fig. 4).

Regarding the linguistic features, we considered four different ones: the inverted frequency of a word irrespective of its context, the surprisal of a word in its context, the associated precision, and the interaction of the surprisal and the precision. We found that the inverted word frequency had a small but significant impact on the neural response: words with a higher frequency (i.e. probability out of context) led to a larger response. Because listeners are exposed to more common words more often, this modulation may emerge due to the long-term plasticity. Similar modulation has been observed before in FFR, where the strength of the response was strongly modulated by the language experience or musical training (Krishnan et al. (2010); Bidelman et al. (2011); Krizman and Kraus (2019)).

Importantly, this effect was present only for the neural response to the high-frequency envelope modulation, but not for that to the fundamental frequency. Because, as discussed above, the former response may contain more cortical contributions than the latter response, the modulation of the neural response by the word frequency may emerge from a cortical rather than subcortical origin. The remaining context-sensitive word-level features did not yield a significant modulation of the neural response at the fundamental frequency. If such a modulation existed, its magnitude were accordingly too weak to be detected in the non-invasive EEG recording.

In summary, we found that the early neural response at the fundamental frequency of speech is predominantly modulated by acoustic features, but also by a linguistic feature, the frequency of a word. The latter result suggests that linguistic processing at the word level involves feedback from higher cortical areas to either very early cortical responses or even further to subcortical structures. We expect that the further investigation of the underlying neural mechanisms will increasingly clarify the role and importance of feedback loops in spoken language processing, with potential applications in speech-recognition technology.

## AUTHOR CONTRIBUTIONS

MK and TR contributed to the conception and design of the study. HW collected and organized the data. MK designed and performed the data analysis. MK wrote the first draft of the manuscript. All authors contributed to manuscript revision, read, and approved the submitted version.

## ACKNOWLEDGMENTS

We would like to thank Dr Katerina Danae Kandylaki for her help with the experimental design and data collection. We are grateful to the Imperial College High Performance Computing Service (doi: 10.14469/hpc/2232).

## DATA AVAILABILITY STATEMENT

EEG data, audio recordings and the word-level features used in this study are available at https://figshare.com/articles/dataset/EEG_recordings_and_stimuli/9033983/1 (Weissbart et al. (2020)). Custom-written Python scripts for computing forward and backward models from the EEG data and the fundamental waveform are available at https://github.com/MKegler/cTRG_toolbox (Etard et al. (2019)).

## Footnotes

1 https://librivox.org/international-short-stories-vol-2-by-william-patten/

2 http://www.gutenberg.org/ebooks/32846

3 We used the open-source Python implementation of the model available at https://github.com/MKegler/pyNSL

4 10.6084/m9.figshare.9033983.v1

